# Crystal structure of SUN1-KASH6 reveals an asymmetric LINC complex architecture compatible with nuclear membrane insertion

**DOI:** 10.1101/2023.06.05.543797

**Authors:** Manickam Gurusaran, Benedikte S. Erlandsen, Owen R. Davies

## Abstract

The LINC complex transmits cytoskeletal forces into the nucleus to control the structure and movement of nuclear contents. It is formed of nuclear SUN and cytoplasmic KASH proteins, which interact within the nuclear lumen, immediately below the outer nuclear membrane. However, the symmetrical location of KASH molecules within SUN-KASH complexes in previous crystal structures has been difficult to reconcile with the steric requirements for insertion of their immediately upstream transmembrane helices into the outer nuclear membrane. Here, we report the crystal structure of the SUN-KASH complex between SUN1 and JAW1/LRMP (KASH6) in an asymmetric 9:6 configuration. This intertwined assembly involves two distinct KASH conformations such that all six KASH molecules emerge on the same molecular surface. Hence, they are ideally positioned for insertion of upstream sequences into the outer nuclear membrane. Thus, we report the first structure of a SUN-KASH complex that is directly compatible with its biological role.

## Introduction

The Linker of Nucleoskeleton and Cytoskeleton (LINC) complex crosses both inner and outer nuclear membranes of the nuclear envelope to mechanically transduce cytoskeletal forces to nuclear contents (Meinke and Schirmer, 2015, Starr and Fridolfsson, 2010, Jahed et al., 2021). It is formed of nuclear SUN (Sad1 and UNC84 homology) and cytoplasmic KASH (Klarsicht, ANC-1, and Syne homology) proteins, which interact within the nuclear lumen (Meinke and Schirmer, 2015, Starr and Fridolfsson, 2010, Jahed et al., 2021). In mammals, there are five SUN proteins (SUN1-5) and multiple isoforms of six KASH proteins (Nesprin-1-4, KASH5 and JAW1/LRMP). These combine to perform the widespread roles of the LINC complex in nuclear structure, shape and positioning (Alam et al., 2015, Luxton et al., 2010, Crisp et al., 2006), in addition to specialised roles in functions such as sound perception in the inner ear and chromosome movements in meiosis (Horn et al., 2013a, Horn et al., 2013b, Lee et al., 2015, Roux et al., 2009). Thus, the LINC complex is necessary for the maintenance of nuclear structure in normal cellular life, in addition to hearing and fertility. Further, LINC complex mutations in humans have been implicated in laminopathies, including Hutchison-Gilford progeria syndrome and Emery-Dreifuss muscular dystrophy (Meinke et al., 2011, Mejat and Misteli, 2010).

The physical interaction between SUN and KASH proteins occurs immediately below the outer nuclear membrane, within the nuclear lumen. This is mediated by the globular SUN domain at the C-terminus of the SUN protein, and the approximately thirty amino-acid KASH domain at the C-terminal end of the KASH protein (Sosa et al., 2012). The widespread functions of SUN proteins are performed by generally expressed, and partially redundant, SUN1 and SUN2 (Lei et al., 2009, Zhang et al., 2009). Their upstream sequences form trimers and higher-order structures (Jahed et al., 2018, Nie et al., 2016, Lu et al., 2008, Hennen et al., 2018), which traverse the nuclear lumen, cross the inner nuclear membrane, and interact with nuclear contents, including the nuclear lamina, chromatin and the telomeric ends of meiotic chromosomes (Crisp et al., 2006, Haque et al., 2006, Haque et al., 2010, Chi et al., 2007, Shibuya et al., 2014). In KASH proteins, transmembrane helices (that cross the outer nuclear membrane) immediately precede the SUN interaction, and lead to diverse cytoplasmic N-terminal domains. The spectrin-repeat domains of widely expressed Nesprin-1 and Nesprin-2 bind to actin (Banerjee et al., 2014, Sakamoto et al., 2017, Zhou et al., 2018). Nesprin-3 interacts via plectin with intermediate filaments (Wilhelmsen et al., 2005). Nesprin-4 functions in sound perception in the inner ear by interacting via kinesin with microtubules (Horn et al., 2013a, Roux et al., 2009). KASH5 functions in meiotic chromosome movements by acting as a transmembrane dynein activating adapter of microtubules (Horn et al., 2013b, Morimoto et al., 2012, Agrawal et al., 2022, Garner et al., 2023). JAW1/LRMP (lymphoid restricted membrane protein) is an atypical KASH protein that is localised in the endoplasmic reticulum and outer nuclear membrane, and is specifically expressed in certain cells of the immune system, small intestine, pancreas and taste buds (Kozono et al., 2018, Okumura et al., 2023, Behrens et al., 1994). It interacts with SUN1 and microtubules, and is required to maintain nuclear shape and Golgi structure (Kozono et al., 2018, Okumura et al., 2023).

The architecture of the LINC complex underpins its mechanism of force transduction. At the heart of this is the structure of the SUN-KASH complex, which lies immediately below the outer nuclear membrane. Crystal structures of SUN-KASH complexes have demonstrated that globular SUN domains form ‘three-leaf clover’-like structures that emanate from upstream trimeric coiled-coils (Sosa et al., 2012, Wang et al., 2012, Zhou et al., 2012). KASH domains are bound at the interface between adjacent protomers, with their C-termini inserted into a KASH-binding pocket and their N-termini interacting with KASH-lids, which are flexible beta hairpin loops within the SUN domain (Sosa et al., 2012, Wang et al., 2012). On the basis of early structures using Nesprin-1/2 sequences, it was assumed that SUN-KASH complexes have a 3:3 stoichiometry (Sosa et al., 2012, Wang et al., 2012). This agreed with the trimeric nature of the luminal regions of SUN proteins, and provided a simple model, in which a linear LINC complex traverses the nuclear envelope and nuclear luminal to transmit forces from the cytoskeleton (Sosa et al., 2013).

In recent crystallographic studies, it was found that non-canonical KASH domains from Nesprin-4 and KASH5 form 6:6 SUN-KASH complexes, in which two 3:3 SUN-KASH trimers interact head-to-head (Gurusaran and Davies, 2021, Cruz et al., 2020). These head-to-head interactions are mediated by zinc-coordination between opposing KASH domains of the Nesprin-4 structure, and through extensive protein-protein interactions between opposing KASH domains of the KASH5 structure (Gurusaran and Davies, 2021). Importantly, it was demonstrated that these 6:6 ‘head-to-head’ complexes represent the solution states of SUN-KASH complexes with Nesprin-4 and KASH5 (Gurusaran and Davies, 2021). In the original SUN-KASH structures with Nesprin-1/2, similar head-to-head interactions between 3:3 complexes were present in the crystal lattice, but were assumed to be crystal contacts (Sosa et al., 2012, Wang et al., 2012). However, SUN1-KASH1 was found to exist solely as a 6:6 complex in solution, and a point mutation targeting the head-to-head interaction, but not affecting constituent 3:3 structures, completely abrogated complex formation *in vitro* (Gurusaran and Davies, 2021). Hence, existing biochemical data indicate that the SUN-KASH complex exists in a 6:6 head-to-head configuration that is essential for its stability.

Whilst the SUN-KASH 6:6 head-to-head complex is consistent with existing biochemical data, little sequence is present between KASH domains and their upstream transmembrane helices. Hence, it is difficult to reconcile how it is sterically possible for all six KASH proteins of a symmetric SUN-KASH 6:6 complex to insert into the outer nuclear membrane (Figure 1a) (Jahed et al., 2021). There are currently two hypotheses for how this may occur *in vivo*. Firstly, that SUN-KASH is disrupted into 3:3 complexes that are oriented vertically such that bound KASH proteins are suitably positioned for insertion of their transmembrane helices into the outer nuclear membrane (Figure 1b) (Sosa et al., 2012, Wang et al., 2012). This is a simple and appealing model, but contradicts existing biochemical data as the SUN-KASH complex has been shown to dissociate in solution upon disruption of the 6:6 interface (Gurusaran and Davies, 2021). Further, this model suggests that cytoplasmic KASH proteins must be either monomers or trimers, whereas KASH5 has been shown to be dimeric (Agrawal et al., 2022, Garner et al., 2023). The second hypothesis is that SUN-KASH 6:6 complexes undergo a hinging motion from their head-to-head configuration, such that all six bound KASH proteins are suitably positioned for membrane insertion (Figure 1c) (Gurusaran and Davies, 2021). This model is supported by computational and biophysical data indicating that such motions are possible and occur in solution (Gurusaran and Davies, 2021), and can explain cytoplasmic KASH proteins forming monomers, dimers or trimers. However, we lack compelling structural evidence in favour of either model.

**Figure 1:**
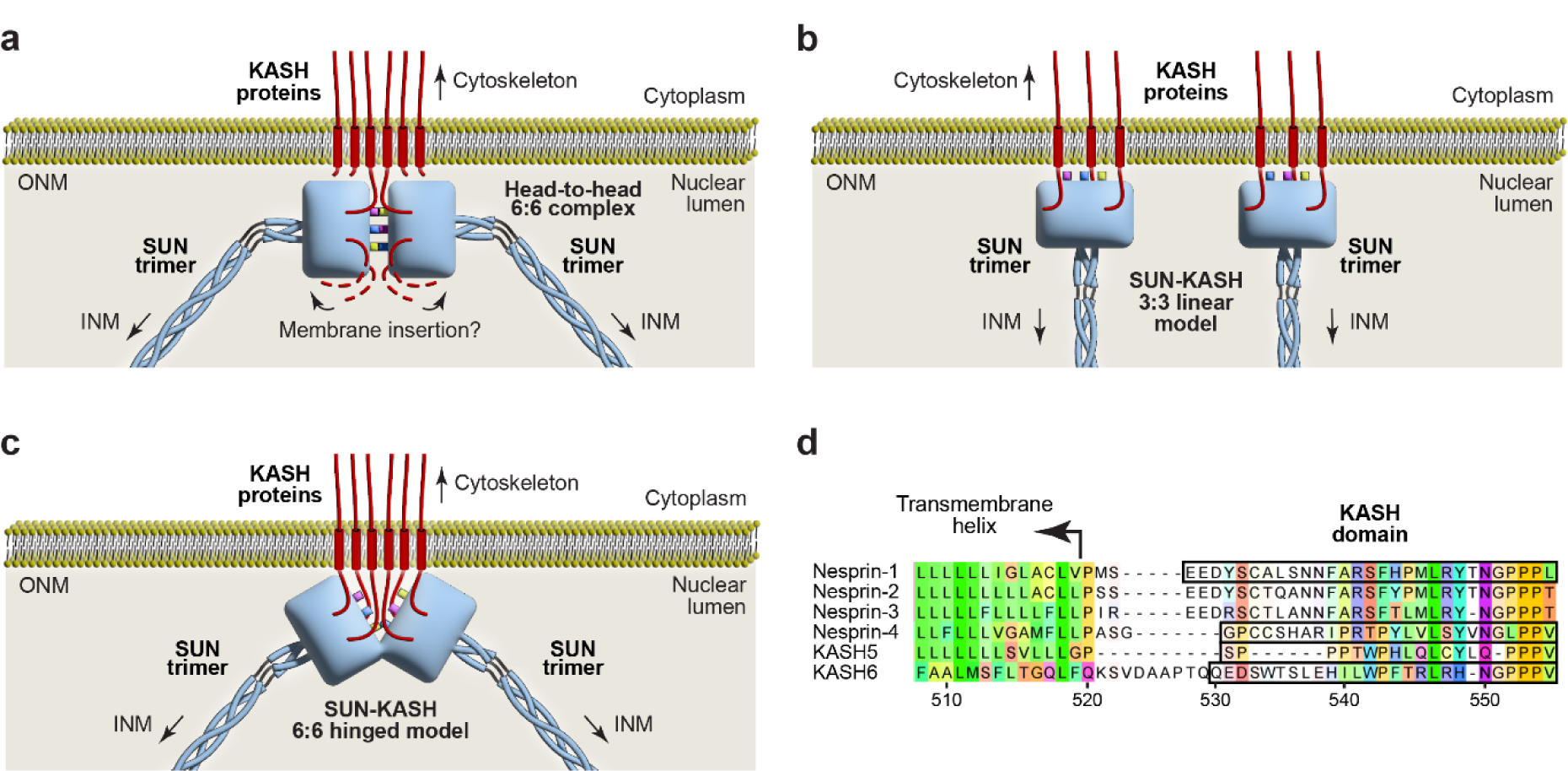
Models of the LINC complex. **(a)** Schematic highlighting the steric challenge of inserting transmembrane helices of all six KASH molecules into the outer nuclear membrane (ONM) from the symmetric 6:6 hetero-oligomer observed in crystal structures of the SUN-KASH complex. (**b,c**) Proposed models of LINC complex structure *in vivo*. (**b**) Linear model in which SUN-KASH 6:6 complexes are disrupted into 3:3 structures that are oriented vertically in the nuclear lumen (Sosa et al., 2012, Wang et al., 2012). (**c**) Hinged model in which SUN-KASH 6:6 complexes are angled to facilitate insertion of their bound KASH molecules into the ONM (Gurusaran and Davies, 2021). (**d**) Structure-based alignment of KASH domains from human KASH proteins in which sequences observed in their crystal structures are highlighted.

Here, we report the crystal structure of the SUN-KASH complex between SUN1 and the divergent KASH domain of JAW1/LRMP (Figure 1d). This reveals an asymmetric 9:6 structure in which three SUN1 trimers are arranged in a triangular configuration, with two distinct KASH peptides bound to each SUN1 trimer. The KASH peptides adopt conformations that are distinct from previous structures, and are arranged such that the N-termini of all six peptides are positioned on the top surface of the molecule. This configuration appears to be compatible with the insertion of upstream KASH sequences into the outer nuclear membrane. We find that the same structure is formed in solution upon transient treatment of the 6:6 complex in mildly acidic conditions. Hence, we report a novel SUN-KASH complex crystal structure that is directly compatible with the steric requirements for insertion of immediately upstream KASH sequences into the outer nuclear membrane.

## Results

### SUN1-KASH6 forms an asymmetric ‘trimer-of-trimers’ structure

In previous SUN-KASH crystal structures, a conserved mechanism of KASH-binding has been observed in which the C-terminus of the KASH domain is inserted into a binding pocket at the interface between adjacent SUN protomers (Sosa et al., 2012, Wang et al., 2012, Cruz et al., 2020, Gurusaran and Davies, 2021). However, divergent sequences of the upstream KASH domain interact differently with the associated KASH-lid of the SUN domain, and have different mechanisms of assembly (Sosa et al., 2012, Wang et al., 2012, Cruz et al., 2020, Gurusaran and Davies, 2021). Indeed, the head-to-head interface between SUN trimers is mediated solely by KASH-lids in Nesprin-1/2 structures, by zinc-coordination between opposing KASH domains in KASH4 structures, and by intertwined interactions between opposing KASH domains and KASH-lids of KASH5 structures (Sosa et al., 2012, Wang et al., 2012, Cruz et al., 2020, Gurusaran and Davies, 2021).

The recently identified KASH domain of JAW1/LRMP (herein referred to as KASH6) diverges from other KASH proteins (Figure 1d), so we wondered whether it may adopt a distinct interaction and assembly mode to those observed in previous structures. Thus, we crystallised the SUN1-KASH6 complex and solved its X-ray crystal structures from several crystals at resolutions between 1.7-2.3 Å (Table 1 and Supplementary Figure 1). In contrast with previous SUN-KASH 6:6 structures, the SUN1-KASH6 crystal structure contains nine SUN domains, such that head-to-head interfaces of three SUN1 trimers interact together around a three-fold symmetry axis in a ‘trimer-of-trimers’ configuration (Figure 2a,b). Hence, the SUN1-KASH6 crystal structure defines a novel LINC complex architecture.

**Figure 2:**
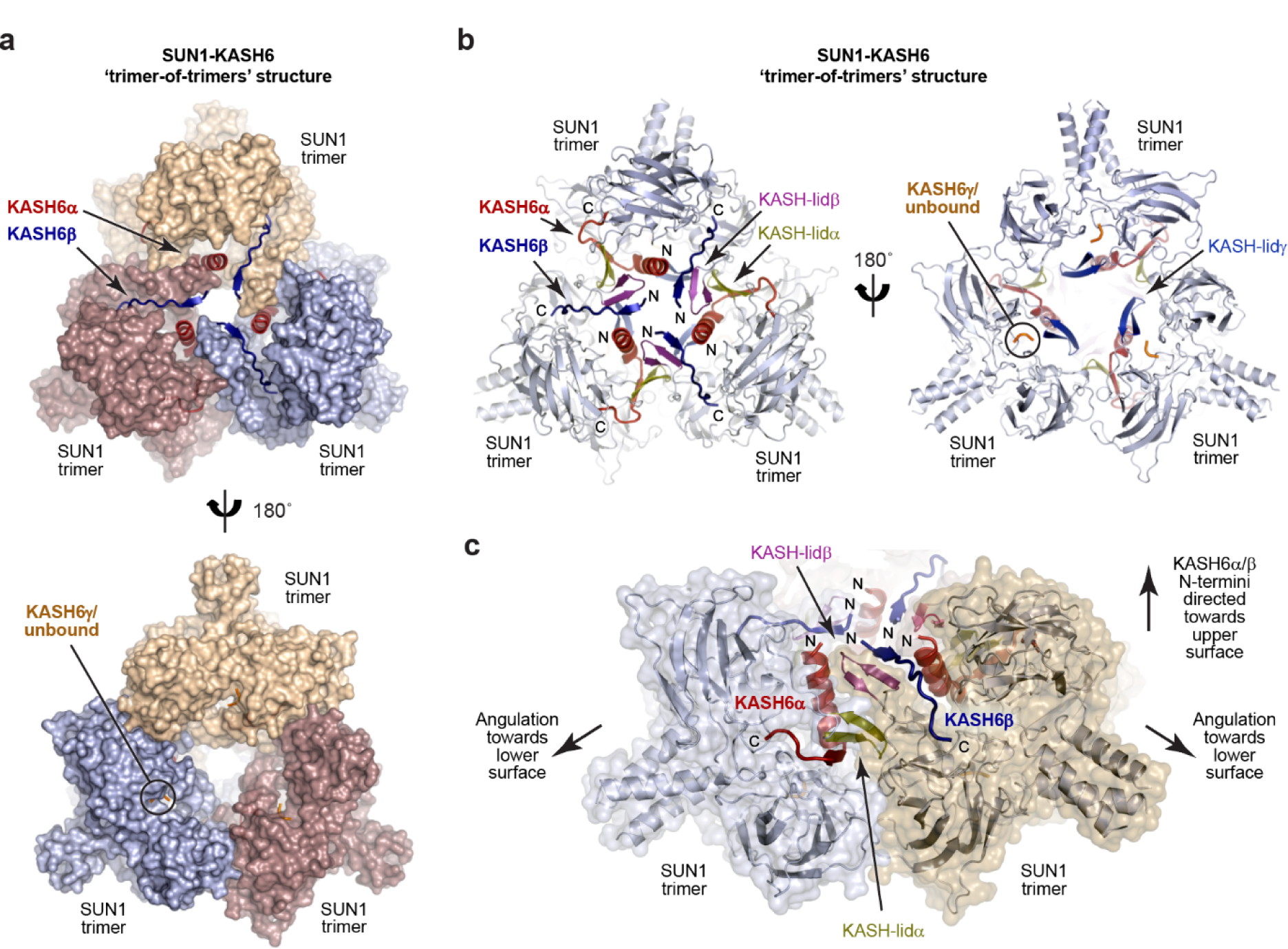
Crystal structure of SUN1-KASH6. (**a-c**) Crystal structure of SUN1-KASH6 (1.7 Å resolution) in a ‘trimer-of-trimers’ configuration. (**a,b**) Overall structure with SUN domains shown as (**a**) surfaces and (**b**) cartoons. The KASH6α and KASH6β molecules bound to the three SUN1 trimers interact extensive with each other, and their associated KASH-lids at the centre of the structure, and emerge on the upper molecular surface. (**c**) The trimer-of-trimers interface is mediated by interactions between KASH6α and KASH6β molecules, and their associated KASH-lids, from adjacent SUN1 trimers. In this interface, KASH-lidα and KASH-lidβ are sandwiched between KASH6α and KASH6β in a six-stranded β-sheet. The N-termini of KASH6α and KASH6β are directed towards the upper surface of the molecule, whilst SUN1 trimers are angled with their N-terminal trimer coiled-coiled pointing downwards towards the lower surface of the molecule.

**Table 1.**
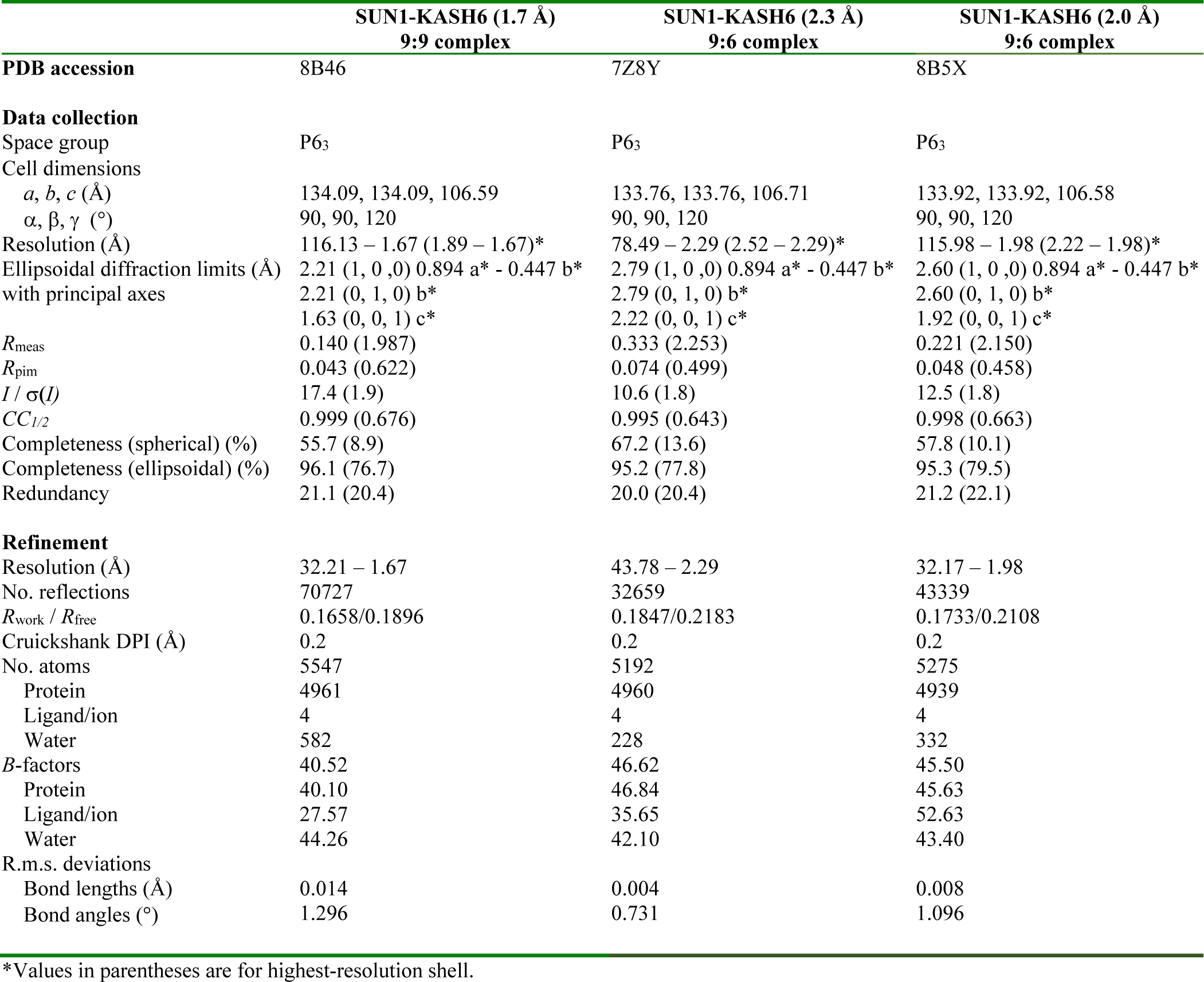
Data collection, phasing and refinement statistics

|  | SUN1-KASH6 (1.7 Å)<br>9:9 complex | SUN1-KASH6 (2.3 Å)<br>9:6 complex | SUN1-KASH6 (2.0 Å)<br>9:6 complex |
| --- | --- | --- | --- |
| **PDB accession** | 8B46 | 7Z8Y | 8B5X |
| **Data collection** |  |  |  |
| Space group | P6<sub>3</sub> | P6<sub>3</sub> | P6<sub>3</sub> |
| Cell dimensions |  |  |  |
| *a*, *b*, *c* (Å) | 134.09, 134.09, 106.59 | 133.76, 133.76, 106.71 | 133.92, 133.92, 106.58 |
| $\alpha$, $\beta$, $\gamma$ (°) | 90, 90, 120 | 90, 90, 120 | 90, 90, 120 |
| Resolution (Å) | 116.13 – 1.67 (1.89 – 1.67)* | 78.49 – 2.29 (2.52 – 2.29)* | 115.98 – 1.98 (2.22 – 1.98)* |
| Ellipsoidal diffraction limits (Å)<br>with principal axes | 2.21 (1, 0, 0) 0.894 *a* * - 0.447 *b* *<br>2.21 (0, 1, 0) *b* *<br>1.63 (0, 0, 1) *c* * | 2.79 (1, 0, 0) 0.894 *a* * - 0.447 *b* *<br>2.79 (0, 1, 0) *b* *<br>2.22 (0, 0, 1) *c* * | 2.60 (1, 0, 0) 0.894 *a* * - 0.447 *b* *<br>2.60 (0, 1, 0) *b* *<br>1.92 (0, 0, 1) *c* * |
| *R*<sub>meas</sub> | 0.140 (1.987) | 0.333 (2.253) | 0.221 (2.150) |
| *R*<sub>pim</sub> | 0.043 (0.622) | 0.074 (0.499) | 0.048 (0.458) |
| *I* / $\sigma(I)$ | 17.4 (1.9) | 10.6 (1.8) | 12.5 (1.8) |
| *CC*<sub>1/2</sub> | 0.999 (0.676) | 0.995 (0.643) | 0.998 (0.663) |
| Completeness (spherical) (%) | 55.7 (8.9) | 67.2 (13.6) | 57.8 (10.1) |
| Completeness (ellipsoidal) (%) | 96.1 (76.7) | 95.2 (77.8) | 95.3 (79.5) |
| Redundancy | 21.1 (20.4) | 20.0 (20.4) | 21.2 (22.1) |
| **Refinement** |  |  |  |
| Resolution (Å) | 32.21 – 1.67 | 43.78 – 2.29 | 32.17 – 1.98 |
| No. reflections | 70727 | 32659 | 43339 |
| *R*<sub>work</sub> / *R*<sub>free</sub> | 0.1658/0.1896 | 0.1847/0.2183 | 0.1733/0.2108 |
| Cruickshank DPI (Å) | 0.2 | 0.2 | 0.2 |
| No. atoms | 5547 | 5192 | 5275 |
| Protein | 4961 | 4960 | 4939 |
| Ligand/ion | 4 | 4 | 4 |
| Water | 582 | 228 | 332 |
| *B*-factors | 40.52 | 46.62 | 45.50 |
| Protein | 40.10 | 46.84 | 45.63 |
| Ligand/ion | 27.57 | 35.65 | 52.63 |
| Water | 44.26 | 42.10 | 43.40 |
| R.m.s. deviations |  |  |  |
| Bond lengths (Å) | 0.014 | 0.004 | 0.008 |
| Bond angles (°) | 1.296 | 0.731 | 1.096 |

\*Values in parentheses are for highest-resolution shell.

Within the trimer-of-trimers structure, SUN1 trimers are positioned with two of their KASH-binding pockets on the sides of the molecule, located at the inter-trimer interface with neighbouring trimers (Figure 2a-c). KASH6 peptides are clearly bound at both of these KASH-binding pockets of each SUN1 trimer (these are herein referred to as KASH6α and KASH6β). The third KASH-binding pocket of each SUN1 trimer is located on the underside of the trimer-of-trimers structure, away from inter-trimer interfaces (Figure 2a,b). Here, we observed poor peptide electron density in the 1.7 Å resolution structure, and could build only three C-terminal KASH6 amino-acids (herein referred to as KASH6γ), with 71% occupancy (Supplementary Figure 2a). In other crystal structures, at resolutions of 2.0 Å and 2.3 Å, we observed no third peptide, with the KASH-binding pocket containing only water molecules (Supplementary Figure 2b). In absence of KASH6γ, the remaining structure was essentially unaltered (r.m.s. deviation = 0.1 Å). Hence, the third binding pocket on the lower surface of the trimer-of-trimers structure cannot stably bind to a KASH6 peptide. We conclude that the SUN1-KASH6 structure is a 9:6 complex in which three SUN1 trimers are bound to six KASH6 peptides.

The KASH6α and KASH6β peptides bound to each SUN1 trimer adopt different conformations. These are arranged such that each inter-trimer interface is formed of intricate interactions between KASH6α and KASH6β peptides, and the associated KASH-lids, of neighbouring SUN1 trimers (Figure 2c). Further, their distinct conformations mean that all KASH6α and KASH6β peptides of the structure are oriented with their N-termini directed towards the top surface of the molecule (Figure 2a-c). This asymmetry within the SUN1-KASH6 trimer-of-trimers structure positions all six stably bound KASH6 peptides in a manner that is suitable for the insertion of their immediately upstream transmembrane helices into the outer nuclear membrane. Hence, uniquely amongst SUN-KASH crystal structures, the architecture of the asymmetric SUN1-KASH6 trimer-of-trimers structure is directly compatible with the steric requirements of membrane insertion, and thereby of LINC complex assembly *in vivo*.

### Distinct KASH domain conformations within the SUN1-KASH6 structure

In previous crystal structures of SUN-KASH complexes, the beta hairpin KASH-lids that extend from SUN domains have retained the same orientation despite differences in the structures adopted by divergent KASH domains (Sosa et al., 2012, Wang et al., 2012, Cruz et al., 2020, Gurusaran and Davies, 2021). In contrast, the distinct KASH6 conformations observed in the SUN1-KASH6 structure are associated with differences in the orientation of their associated KASH-lids (Supplementary Figure 3). The KASH-lid associated with KASH6α adopts the canonical conformation (Figure 3a). In contrast, the KASH-lid associated with KASH6β has a high angulation (Figure 3b), and the KASH-lid at the KASH6γ/unbound site has a low angulation (Figure 3c). These are herein referred to as KASH-lidα, KASH-lidβ and KASH-lidγ, respectively.

**Figure 3:**
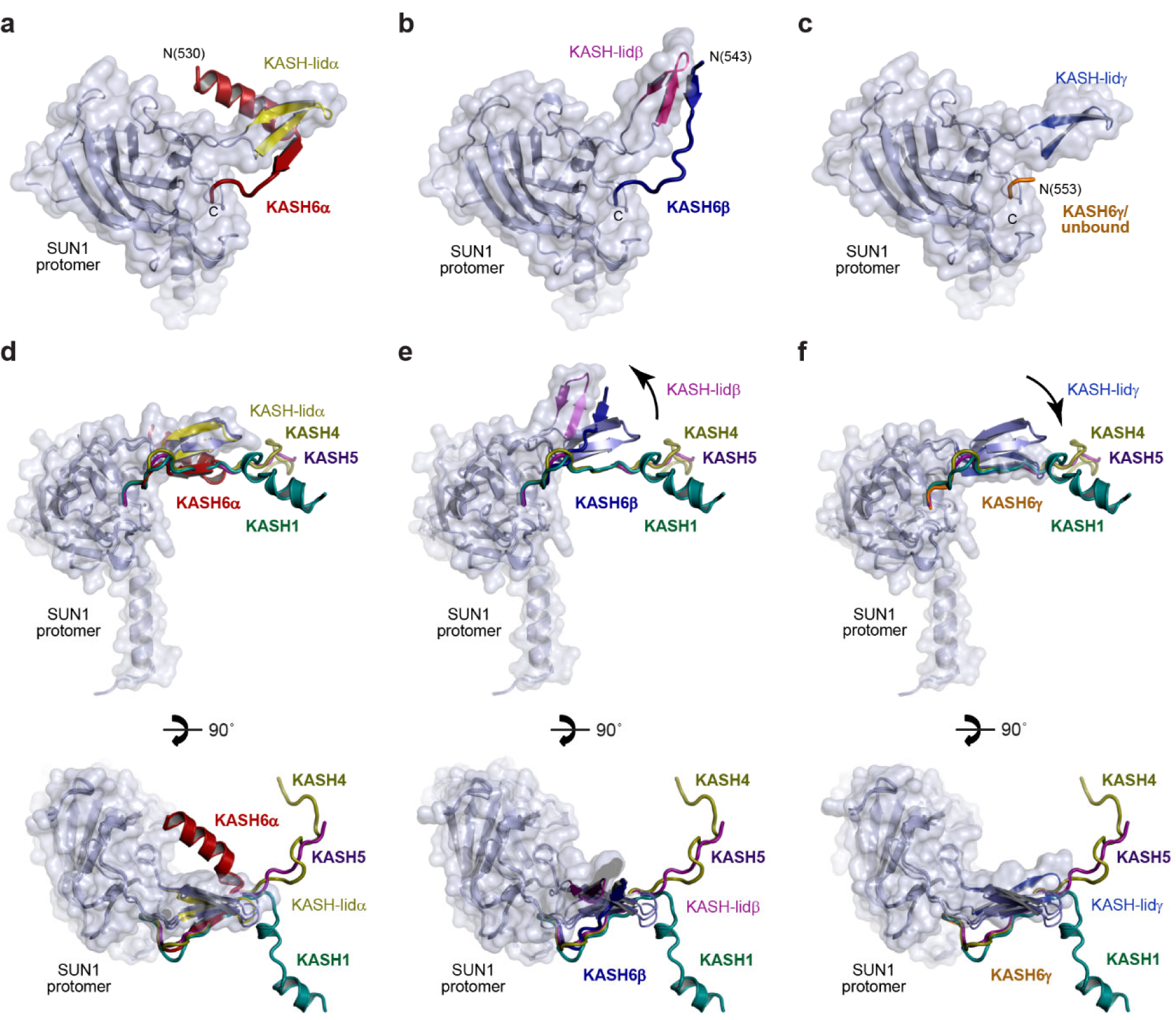
Unique conformations of KASH6 peptides within the SUN1-KASH6 structure. (**a-c**) Alternative conformations adopted by KASH6 peptides and their associated SUN domain KASH-lids for (**a**) KASH6α, (**b**) KASH6β and (**c**) KASH6γ. (**d-f**) Superposition of the KASH peptide and its associated SUN1 protomer from previous crystal structures of SUN1-KASH1 (green; PDB accession 6R15; Gurusaran and Davies, 2021), SUN1-KASH4 (brown; PDB accession 6R16; Gurusaran and Davies, 2021) and SUN1-KASH5 (purple; PDB accession 6R2I; Gurusaran and Davies, 2021), with (**d**) SUN1-KASH6α (red), (**e**) SUN1-KASH6β (blue) and (**f**) SUN1-KASH6γ (yellow). (**d**) KASH6α follows the same initial path, undergoing β-interaction with its canonically-oriented KASH-lid, as in all previous SUN1-KASH structures. It then hooks below KASH-lidα, undergoing a 90° bend to form an α-helix that is oriented in the opposite direction to the partial α-helical end of KASH1. (**e**) KASH6β forms the same β-interaction with its KASH-lid as in all other SUN1-KASH structures, but with an unusually high angulation of both KASH6β and KASH6-lidβ. (**f**) KASH6γ was only observed as three amino-acids at the KASH-binding pocket, with partial occupancy, and in only a subset of crystal structures. In absence of the β-interaction, its KASH-lid adopts a low angulation in which its first β-strand overlays with the KASH-lid-interacting β-strand of other KASH peptides.

The KASH6α peptide follows the canonical binding mode of C-terminal insertion into the KASH-binding pocket, followed by a β-interaction with the canonical KASH-lidα (Figure 3a) (Sosa et al., 2012, Gurusaran and Davies, 2021, Cruz et al., 2020, Wang et al., 2012). However, KASH6α then hooks underneath the KASH-lid, and undergoes a 90° turn to form an N-terminal α-helix that is perpendicular to the coiled-coil axis of the SUN1 trimer (Figure 3a). This conformation resembles that of KASH1/2 structures, although the turn is in the opposite direction, such that N-terminal α-helices of KASH6α point inwards towards the centre of the SUN1 trimer, whereas those of KASH1/2 point outwards (Figure 3d). Importantly, the three KASH6α α-helices of the trimer-of-trimers structure are parallel, with their N-termini directed towards the top surface of the molecule (Figure 2a-c).

The KASH6β peptide also follows the canonical pattern of C-terminal insertion into the KASH-binding pocket and β-interaction with KASH-lidβ (Figure 3b). However, it does not hook under the KASH-lid or form an N-terminal α-helix. Instead, the β-structure formed by KASH6β is extended, in a manner similar to KASH5 (Figure 3b,e) (Gurusaran and Davies, 2021, Cruz et al., 2020). The resultant structure with KASH-lidβ is unusually highly angled, directed away from the core of the SUN1 trimer (Figure 3b). The N-termini of the three KASH6β peptides of the structure point towards the same molecular surface, and are each sandwiched between the N-terminal α-helices of two KASH6α peptides (Figure 2a-c). Hence, the N-termini of all six KASH6 peptides are located in closely proximity, on the upper surface of the molecule.

At the remaining KASH6γ/unbound site, KASH-lidγ adopts a low angled conformation in which its β-structure follows the path that would normally be occupied by the KASH peptide (Figure 3c,f). This localisation likely acts as a steric hindrance for KASH-binding, accounting for the lack of stable binding by a third KASH6 peptide.

### KASH domains and KASH-lids form extensive inter-trimer interfaces

The inter-trimer interfaces that hold together the SUN1-KASH6 structure are formed by the KASH6α and KASH6β molecules of neighbouring SUN1 trimers interacting via their associated KASH-lids (Figure 2c). At its core, the inter-trimer interface is formed of a six-stranded β-sheet, in which KASH-lidα and KASH-lidβ of adjacent SUN1 trimers interact together, sandwiched between the β-strands of their KASH6α and KASH6β peptides (Figure 2c). This interaction mode resembles the KASH-mediated head-to-head interface of the SUN1-KASH5 6:6 structure (Gurusaran and Davies, 2021, Cruz et al., 2020). The inter-trimers interface is stabilised by hydrophobic interactions between the surface of the β-sheet (including SUN1 amino-acids F671, I673, L675 and W676) and the N-terminal α-helix of KASH6α (including KASH6 amino-acids W534, L537 and L541) (Figure 4a). It is further supported by interactions of the unbound KASH-lidγ, including binding of its tip amino-acids F671 and I673 with an opposing KASH-lidα (Figure 4b), in a manner that resembles the head-to-head interface of the SUN1-KASH1 6:6 structure (Gurusaran and Davies, 2021, Cruz et al., 2020). Further, these KASH-lidγ interactions position SUN1 promoters with their coiled-coils angled downwards, away from the upper surface at which the N-termini of KASH6 molecules presented (Figure 2c). Hence, intricate interactions between KASH6 molecules and KASH-lids of adjacent SUN1 trimers form extensive inter-trimer interfaces that establish a trimer-of-trimers platform in which KASH6 molecules are directed to the upper surface and SUN1 coiled-coils are angled towards the lower surface. This architecture appears ideal for its known biological location in the nuclear lumen, with immediately upstream KASH6 sequences inserted into the outer nuclear membrane.

**Figure 4:**
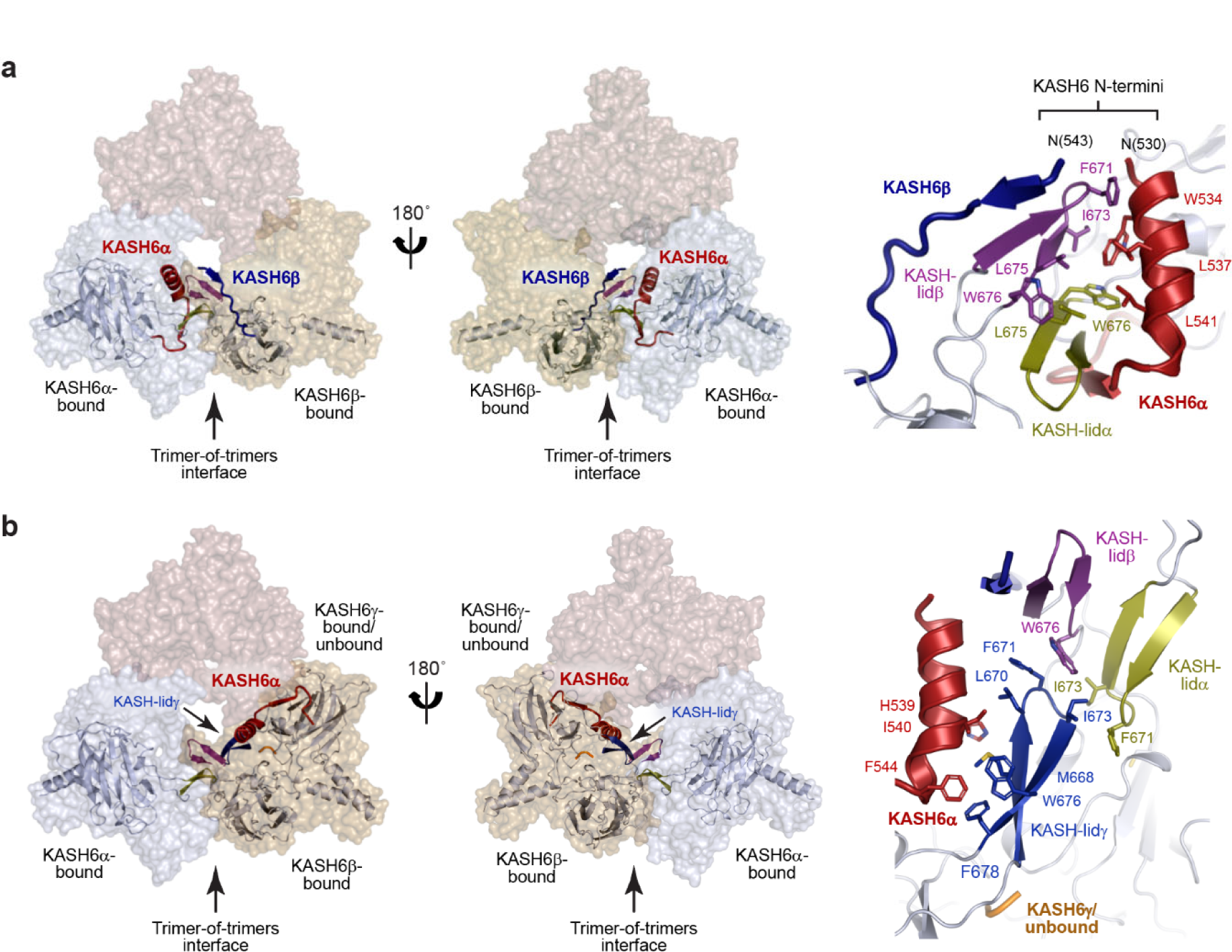
Structural roles of KASH6 peptides and KASH-lids within the trimer-of-trimers assembly. (**a,b**) Location of interacting KASH6 peptides and associated SUN domains within the trimer-of-trimers structure (left), and their molecular details (right). (**a**) The trimer-of-trimers interface is mediated by an interaction between KASH6α (and associated SUN domain KASH-lidα) and KASH6β (and associated SUN domain KASH-lidβ) of adjacent SUN1 trimers. The KASH6α and KASH6β peptides sandwich their KASH-lids together in a β-sheet structure that is stabilised by hydrophobic packing between the KASH6α helix and the β -sheet. This results in the N-termini of KASH6α and KASH6β peptides being oriented close together on the upper molecular surface. (**b**) KASH-lidγ of the KASH6γ-associated or unbound SUN1 protomer interacts with the KASH6α helix and KASH-lidβ of the same SUN1 trimer, and KASH-lidα of the adjacent SUN1 trimer. Its interaction with KASH-lidβ involves the same molecular contacts as the head-to-head interfaces of SUN1-KASH1 and SUN1-KASH5 6:6 complexes (Gurusaran and Davies, 2021).

### SUN1-KASH6 undergoes trimer-of-trimers assembly in solution

Does SUN1-KASH6 form the same trimer-of-trimers structure in solution? We utilised size-exclusion chromatography multi-angle light scattering (SEC-MALS) to determine its solution oligomeric state. SEC-MALS analysis revealed that SUN1-KASH6 exists predominantly as a 6:6 complex (150 kDa) (Figure 5a), similar to other SUN-KASH complexes (Gurusaran and Davies, 2021). As all diffracting crystals were obtained in acidic conditions, we wondered whether trimer-of-trimers formation may occur at low pH. Hence, we treated SUN1-KASH6 in mildly acidic (pH 5.0) conditions. Upon subsequent SEC-MALS analysis at pH 7.5, we observed the presence of roughly equal amounts of a 9:6 complex (212 kDa) and a 6:6 complex (150 kDa) (Figure 5a). This transition became almost complete when we truncated the N-terminal end of the KASH6 peptide to match only the amino-acids observed in the crystal structure (Figure 5b). Hence, SUN1-KASH6 underwent irreversible conformational change from a 6:6 complex to a 9:6 complex upon transient treatment in mildly acidic conditions in solution.

**Figure 5:**
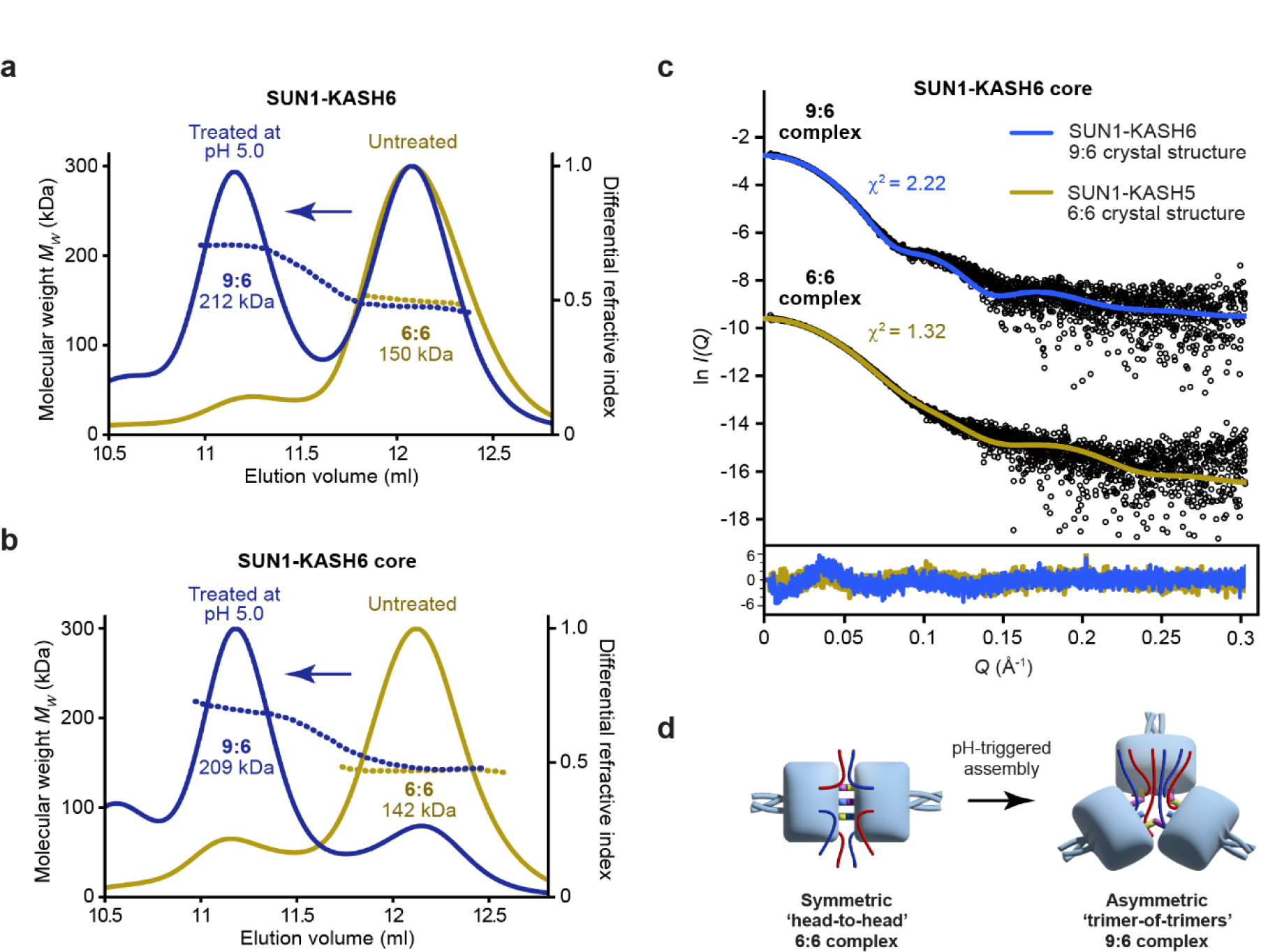
Assembly of SUN1-KASH6 9:6 complexes in solution. (**a,b**) Size-exclusion chromatography multi-angle light scattering showing that (**a**) SUN1-KASH6 and (**b**) SUN1-KASH6 core are 6:6 complexes that assemble into 9:6 complexes following treatment at pH 5.0 (theoretical 9:6 – 232 kDa and 222 kDa; theoretical 6:6 – 165 kDa and 155 kDa). (**c**) Small-angle X-ray scattering of SUN1-KASH6 core 9:6 and 6:6 species overlaid with the theoretical scattering curves of the 9:6 and 6:6 crystal structures of SUN1-KASH6 and SUN1-KASH5 (PDB accession 6R2I; Gurusaran and Davies, 2021), showing χ^2^ values of 2.22 and 1.32, respectively (for other combinations, χ^2^ > 24). (**d**) SUN1-KASH6 forms a symmetric head-to-head 6:6 complex in solution that undergoes pH-triggered conformational change into an asymmetric trimer-of-trimers 9:6 complex.

We next assessed the solution structure of the SUN1-KASH6 9:6 and 6:6 complexes by size-exclusion chromatography small angle X-ray scattering. The SAXS scattering curve of the 9:6 complex was closely fitted by the SUN1-KASH6 trimer-of-trimers 9:6 crystal structure (χ^2^ = 2.22) (Figure 5c and Supplementary Figure 4a,b), but not the previous SUN1-KASH5 head-to-head 6:6 crystal structure (χ^2^ = 62.5). In contrast, the SAXS scattering curve of the 6:6 complex was closely fitted by the SUN1-KASH5 head-to-head 6:6 crystal structure (χ^2^ = 1.32) (Figure 5c and Supplementary Figure 4a,b), but not the SUN1-KASH6 trimer-of-trimers 9:6 crystal structure (χ^2^ = 24.5). These data strongly support that the SUN1-KASH6 9:6 complex in solution has the same trimer-of-trimers configuration that is observed in the crystal structure, and that the 6:6 complex in solution has the same head-to-head structure that was previously observed for other SUN-KASH complexes.

In summary, we find that SUN1-KASH6 forms a symmetric head-to-head 6:6 structure that undergoes irreversible conformational change to an asymmetric trimer-of-trimers 9:6 structure upon transient treatment in mildly acidic conditions (Figure 5d). This conformational change requires the dissociation of two KASH6 molecules from each initial 6:6 complex. Hence, the few amino-acids of a KASH6γ peptide observed with partial occupancy in the third KASH-binding pocket of the 1.7 Å crystal structure were likely remnants of the peptide having failed to fully dissociate upon trimer-of-timers assembly. Importantly, the pH-triggered 9:6 assembly constitutes the first SUN-KASH complex structure in which all bound KASH peptides are positioned on the top surface of the molecule. Thus, we report a novel SUN-KASH complex crystal structure that is also formed in solution, which is directly compatible with the steric requirements for insertion of upstream transmembrane helices of all bound KASH molecules into the outer nuclear membrane (Figure 6).

**Figure 6:**
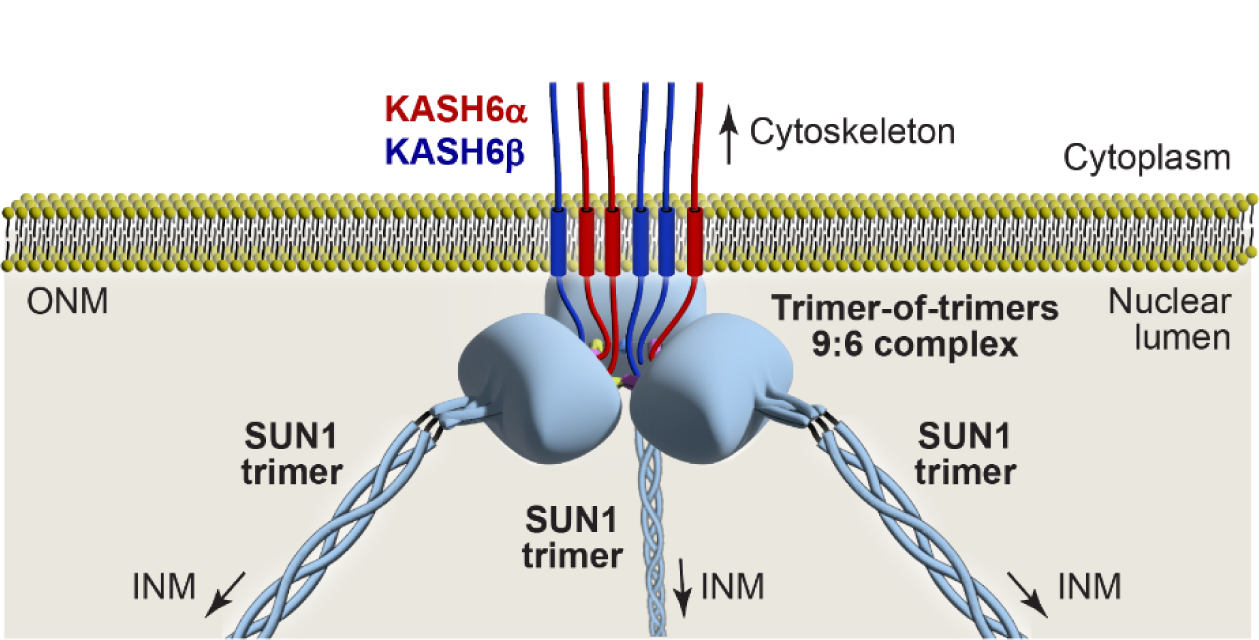
Model of a SUN1-KASH6 9:6 LINC complex. Schematic model of how a SUN1-KASH6 asymmetric trimer-of-trimers LINC complex may be positioned with the nuclear lumen. The six bound KASH6 domains emerge from the 9:6 complex on the top surface of the molecule, favouring of their immediately upstream transmembrane helices into the outer nuclear membrane (ONM). The SUN1 trimers are tilted downwards within the trimer-of-trimers structure, facilitating the angled passage of SUN1’s upstream trimeric coiled-coil across the nuclear lumen, towards the inner nuclear membrane (INM).

## Discussion

The SUN1-KASH6 9:6 crystal structure provides a new snapshot of the molecular architecture that may be adopted by a LINC complex. In this structure, SUN1 trimers are arranged around a three-fold symmetry axis, such that their upstream trimeric coiled-coils emanate at the points of a triangle. In keeping with previous SUN-KASH structures, KASH6 C-termini are bound in pockets between adjacent SUN1 protomers (Sosa et al., 2012, Wang et al., 2012, Cruz et al., 2020, Gurusaran and Davies, 2021). However, whilst the two binding pockets at the inter-trimer interface were occupied with KASH6 peptides, the third pocket on the lower surface of the molecule was either unoccupied or contained only a few C-terminal amino-acids. Hence, the third pocket does not stably bind KASH6 peptides. The two robustly bound KASH6 peptides of each SUN1 trimer adopt distinct conformations from each other, and from other KASH domains (Sosa et al., 2012, Wang et al., 2012, Cruz et al., 2020, Gurusaran and Davies, 2021). KASH6α undergoes a β-interaction with its associated KASH-lid, which it hooks underneath, and then forms an α-helix perpendicular to the SUN1 trimer. In contrast, KASH6β forms a more extensive β-interaction with its KASH-lid. At each inter-trimer interface, KASH6α and KASH6β from interacting SUN1 trimers form integrated structures, in which their associated KASH-lids are sandwiched within a six-stranded β-sheet. Hence, this highly intertwined structure is stabilised by extensive interactions between adjacent SUN1 trimers and KASH peptides. Importantly, the distinct KASH6α and KASH6β conformations result in an asymmetrical structure in which the N-termini of all six KASH6 peptides emanate towards the top surface of the molecule. This configuration may favour insertion of upstream sequences of all bound KASH6 molecules into the outer nuclear membrane. Hence, uniquely amongst SUN-KASH crystal structures, the SUN1-KASH6 structure appears to be directly compatible with outer nuclear membrane insertion (Figure 6).

In solution, SUN1-KASH6 adopted the same 6:6 complex as other SUN-KASH complexes (Gurusaran and Davies, 2021), but formed the 9:6 complex observed in the crystal structure upon treatment in mildly acidic conditions. How does this relate to the likely mechanism of SUN-KASH6 assembly *in vivo*? Our findings may reflect a true requirement for protonation *in vivo*, which could be achieved through a high local proton concentration, an interacting partner or during the folding process. However, an alternative explanation is that low pH simply overcame an energy barrier between conformations that enabled formation of the 9:6 species. This hypothesis is supported by our observation that following transient treatment at pH 5, the assembled 9:6 complex subsequently remained stable at pH 7.5. Hence, the energy barrier between conformations may be overcome by other factors *in vivo*, such as the process of membrane insertion, and the steric and chemical effects of membrane proximity.

What is the structure of the LINC complex *in vivo*? The only reliable answer to this question will come from its direct visualisation *in situ* by cryo-electron microscopy. However, current technologies are not yet sufficient to resolve narrow coiled-coils and small complexes associated with membrane surfaces within cells. Hence, we are currently limited to inferring its possible cellular architecture based on crystal structures and *in vitro* biophysical data. As discussed, the SUN1-KASH6 9:6 structure represents an elaborate assembly of SUN1 and KASH6 molecules in a manner that is clearly compatible with outer nuclear membrane insertion, so it seems likely that this structure forms *in vivo* (Figure 6). We did not observe 9:6 assembly upon acidic treatment of other LINC complexes (SUN1-KASH1, SUN1-KASH4 and SUN1-KASH5), so our observations of 9:6 assembly may be specific to the divergent KASH6 sequence. Nevertheless, the wider principle of higher order asymmetric assembly, owing to a trigger or the steric effects of membrane proximity/insertion, may be broadly applicable to LINC complexes. Indeed, we previously reported that SUN1-KASH4 formed a 12:12 species in addition to its predominant 6:6 complex (Gurusaran and Davies, 2021). Further, different LINC complex architectures may form in different cells, at different times, and in response to different stresses (Jahed et al., 2021). Hence, to explain the varied and dynamic roles of LINC complexes in cells, we must consider all possible architectures that have been hypothesised and observed, including 3:3 linear complexes (Sosa et al., 2012, Wang et al., 2012), hinged 6:6 complexes (Gurusaran and Davies, 2021), and the newly observed asymmetric 9:6 architecture of SUN1-KASH6.

Overall, the SUN1-KASH6 structure reported herein establishes three novel precedents. Firstly, stable SUN-KASH LINC complexes can include unoccupied KASH-binding pockets. Secondly, the same KASH peptide, and associated KASH-lid, can adopt alternative conformations within the same SUN-KASH complex. Finally, SUN-KASH complexes can form asymmetric assemblies in which all bound KASH peptides emerge on the same surface of the complex. Hence, it is possible for a LINC complex to assemble in which all bound KASH molecules are oriented in a manner suitable for insertion into the outer nuclear membrane (Figure 6). Whilst our findings relate to SUN1-KASH6, it is possible that other LINC complexes may undergo conformational change into similar higher order assemblies for membrane insertion. Indeed, the trimer-of-trimers interface of SUN1-KASH6 involves the hinging motion that we previously observed for other SUN-KASH molecules (Gurusaran and Davies, 2021). Thus, we provide the first reported case in which an experimental structure of a LINC core complex, and its solution biophysical data, are immediately compatible with outer nuclear membrane insertion.

## Materials and Methods

### Recombinant protein expression and purification

The SUN domain sequence of human SUN1 (amino-acids 616-812) was fused to a TEV-cleavable N-terminal His_6_-GCN4 tag and cloned into a pRSF-Duet1 vector (Merck Millipore). The KASH domain sequence of human JAW1/LRMP (amino-acids 515-555 and 531-555 for KASH6 and KASH6 core, respectively) was cloned into a pMAT11 vector (Peranen et al., 1996) for expression with an N-terminal His_6_-MBP tag. SUN1 and KASH6 were co-expressed in BL21 (DE3) cells (Novagen) in 2xYT media, and induced at an OD of 0.8 with 0.5 mM IPTG for 16 hours at 25°C. Cells were lysed by cell disruption in 20 mM Hepes (pH 7.5), 500 mM KCl, and lysate was purified through consecutive Ni-NTA (Qiagen), amylose (NEB) and HiTrap Q HP (GE Healthcare) anion exchange chromatography. Fusion proteins were cleaved by TEV protease, and cleaved samples were purified by HiTrap Q HP anion exchange, followed by size-exclusion chromatography (HiLoad 10/300 Superdex 200, GE Healthcare) in 20 mM Hepes (pH 7.5), 150 mM KCl, 2 mM DTT. Protein samples were concentrated using Amicon Ultra-4 10 MWCO Centrifugal devices (Thermo Scientific), and flash frozen in liquid nitrogen and stored in −80°C. Protein samples were analysed by SDS-PAGE with Coomassie staining, and concentrations were determined by UV spectroscopy using a Cary 60 UV spectrophotometer (Agilent) with extinction coefficients and molecular weights calculated by ProtParam (http://web.expasy.org/protparam/).

### Crystallisation and structure solution of SUN1-KASH6 (1.7 Å) at 9:9 stoichiometry

SUN1-KASH6 protein crystals were obtained through vapour diffusion in sitting drops, by mixing protein at 12 mg/ml with crystallisation solution (0.15 M Magnesium chloride, 0.1 M MES pH 6.0, 3% w/v PEG 6000) and equilibrating at 20°C. Crystals were cryo-protected by addition of 25% ethylene glycol, and were cryo-cooled in liquid nitrogen. X-ray diffraction data were collected at 0.6702 Å, 100 K, as 3600 consecutive 0.10° frames of 0.005 s exposure on a Pilatus3 6M detector at beamline I24 of the Diamond Light Source synchrotron facility (Oxfordshire, UK). Data were processed using *AutoPROC* (Vonrhein et al., 2011), in which indexing, integration, scaling were performed by *XDS* (Kabsch, 2010) and *Aimles s*(Evans, 2011), and anisotropic correction with a local *I/σ(I)* cut-off of 1.2 was performed by *STARANISO* (Tickle et al., 2018). Crystals belong to hexagonal spacegroup P6_3_ (cell dimensions a = 134.09 Å, b = 134.09 Å, c = 106.59 Å, α = 90°, β = 90°, γ = 120°), with one 3:3 SUN1-KASH6 complex in the asymmetric unit. Structure solution was achieved by molecular replacement using *PHASER* (McCoy et al., 2007), in which three copies of the SUN domain from the SUN1-KASH5 structure (PDB accession 6R2I; Gurusaran and Davies, 2021) were placed in the asymmetric unit. Model building was performed through iterative re-building by *PHENIX Autobuild* (Adams et al., 2010) and manual building in *Coot* (Emsley et al., 2010). The structure was refined using *PHENIX refine* (Adams et al., 2010), using isotropic (water) atomic displacement parameters, with 22 TLS groups. The structure was refined against data to anisotropy-corrected data with resolution limits between 1.67 Å and 2.35 Å, to *R* and *R_free_* values of 0.1658 and 0.1896, respectively, with 96.41% of residues within the favoured regions of the Ramachandran plot (0 outliers), clashscore of 7.38 and overall *MolProbity* score of 1.64 (Chen et al., 2010). The final SUN1-KASH6 model was analysed using the Online_DPI webserver (http://cluster.physics.iisc.ernet.in/dpi) to determine a Cruikshank diffraction precision index (DPI) of 0.2 Å (Kumar et al., 2015).

### Crystallisation and structure solution of SUN1-KASH6 (2.3 Å) at 9:6 stoichiometry

SUN1-KASH6 protein crystals were obtained through vapour diffusion in sitting drops, by mixing protein at 12 mg/ml with crystallisation solution (0.1 M Bis Tris pH 5.5, 0.3 M magnesium formate) and equilibrating at 20°C. Crystals were cryo-protected by addition of 30% ethylene glycol, and were cryo-cooled in liquid nitrogen. X-ray diffraction data were collected at 0.9999 Å, 100 K, as 3600 consecutive 0.10° frames of 0.010 s exposure on a Pilatus3 6M detector at beamline I24 of the Diamond Light Source synchrotron facility (Oxfordshire, UK). Data were processed using *AutoPROC* (Vonrhein et al., 2011), in which indexing, integration, scaling were performed by *XDS* (Kabsch, 2010) and *Aimless* (Evans, 2011), and anisotropic correction with a local *I/σ(I)* cut-off of 1.2 was performed by *STARANISO* (Tickle et al., 2018). Crystals belong to hexagonal spacegroup P6_3_ (cell dimensions a = 133.76 Å, b = 133.76 Å, c = 106.71 Å, α = 90°, β = 90°, γ = 120°), with one 3:2 SUN1-KASH6 complex in the asymmetric unit. Structure solution was achieved by molecular replacement using *PHASER* (McCoy et al., 2007), in which three copies of the SUN domain from the SUN1-KASH5 structure (PDB accession 6R2I; Gurusaran and Davies, 2021) were placed in the asymmetric unit. Model building was performed through iterative re-building by *PHENIX Autobuild* (Adams et al., 2010) and manual building in *Coot* (Emsley et al., 2010). The structure was refined using *PHENIX refine* (Adams et al., 2010), using isotropic (water) atomic displacement parameters, with 16 TLS groups. The structure was refined against data to anisotropy-corrected data with resolution limits between 2.29 Å and 2.93 Å, to *R* and *R_free_* values of 0.1847 and 0.2183, respectively, with 97.06% of residues within the favoured regions of the Ramachandran plot (0 outliers), clashscore of 4.92 and overall *MolProbity* score of 1.42 (Chen et al., 2010). The final SUN1-KASH6 model was analysed using the Online_DPI webserver (http://cluster.physics.iisc.ernet.in/dpi) to determine a Cruikshank diffraction precision index (DPI) of 0.2 Å (Kumar et al., 2015).

### Crystallisation and structure solution of SUN1-KASH6 (2.0 Å) at 9:6 stoichiometry

SUN1-KASH6 protein crystals were obtained through vapour diffusion in sitting drops, by mixing protein at 12 mg/ml with crystallisation solution (0.15 M sodium acetate; 0.1 M Sodium citrate tribasic dihydrate pH 5.5) and equilibrating at 20°C. Crystals were cryo-protected by addition of 25% ethylene glycol, and were cryo-cooled in liquid nitrogen. X-ray diffraction data were collected at 0.6702 Å, 100 K, as 3600 consecutive 0.10° frames of 0.005 s exposure on a Pilatus3 6M detector at beamline I24 of the Diamond Light Source synchrotron facility (Oxfordshire, UK). Data were processed using *AutoPROC* (Vonrhein et al., 2011), in which indexing, integration, scaling were performed by *XDS* (Kabsch, 2010) and *Aimless* (Evans, 2011), and anisotropic correction with a local *I/σ(I)* cut-off of 1.2 was performed by *STARANISO* (Tickle et al., 2018). Crystals belong to hexagonal spacegroup P6_3_ (cell dimensions a = 134.09 Å, b = 134.09 Å, c = 106.59 Å, α = 90°, β = 90°, γ = 120°), with one 3:2 SUN1-KASH6 complex in the asymmetric unit. Structure solution was achieved by molecular replacement using *PHASER* (McCoy et al., 2007), in which a 3:2 complex from the SUN1-KASH6 9:9 structure (PDB accession 8B46) was placed in the asymmetric unit. Model building was performed through iterative re-building by *PHENIX Autobuild* (Adams et al., 2010) and manual building in *Coot* (Emsley et al., 2010). The structure was refined using *PHENIX refine* (Adams et al., 2010), using isotropic (water) atomic displacement parameters, with 16 TLS groups. The structure was refined against data to anisotropy-corrected data with resolution limits between 1.98 Å and 2.77 Å, to *R* and *R_free_* values of 0.1733 and 0.2107, respectively, with 96.40% of residues within the favoured regions of the Ramachandran plot (0 outliers), clashscore of 6.39 and overall *MolProbity* score of 1.59 (Chen et al., 2010). The final SUN1-KASH6 model was analysed using the Online_DPI webserver (http://cluster.physics.iisc.ernet.in/dpi) to determine a Cruikshank diffraction precision index (DPI) of 0.2 Å (Kumar et al., 2015).

### Size-exclusion chromatography multi-angle light scattering (SEC-MALS)

The absolute molecular masses of SUN1-KASH6 complexes were determined by size-exclusion chromatography multi-angle light scattering (SEC-MALS). Protein samples at 8-13 mg/ml were loaded onto a Superdex™ 200 Increase 10/300 GL size exclusion chromatography column (GE Healthcare) in 20 mM Hepes pH 7.5, 150 mM KCl, 2 mM DTT, at 0.5 ml/min using an ÄKTA™ Pure (GE Healthcare). For induction of 9:6 complex formation, samples were treated in 20 mM Sodium acetate pH 5.0, 150 mM KCl, 10% glycerol, 2 mM DTT overnight, and were analysed by size-exclusion chromatography in the buffer described above. The column outlet was fed into a DAWN® HELEOS™ II MALS detector (Wyatt Technology), followed by an Optilab® T-rEX™ differential refractometer (Wyatt Technology). Light scattering and differential refractive index data were collected and analysed using *ASTRA*® 6 software (Wyatt Technology). Molecular weights and estimated errors were calculated across eluted peaks by extrapolation from Zimm plots using a *dn/dc* value of 0.1850 ml/g. SEC-MALS data are presented as differential refractive index (dRI) profiles, with fitted molecular weights (*M_W_*) plotted across elution peaks.

### Size-exclusion chromatography small-angle X-ray scattering (SEC-SAXS)

SEC-SAXS experiments were performed at beamline B21 of the Diamond Light Source synchrotron facility (Oxfordshire, UK). Protein samples at 12-25 mg/ml were prepared as described above for SEC-MALS and were loaded onto a Superdex™ 200 Increase 10/300 GL size exclusion chromatography column (GE Healthcare) in 20 mM Hepes pH 7.5, 150 mM KCl, 2 mM DTT, at 0.5 ml/min using an Agilent 1200 HPLC system. The column outlet was fed into the experimental cell, and SAXS data were recorded at 12.4 keV, detector distance 4.014 m, in 3.0 s frames. Data were subtracted, averaged and analysed for Guinier region *Rg* using *ScÅtter* 3.0 (http://www.bioisis.net), and *P(r)* distributions were fitted using *PRIMUS* (P.V.Konarev, 2003). Crystal structures were fitted to experimental data using *CRYSOL* (Svergun D.I., 1995). The SUN1-KASH6 9:6 crystal structure fitted to 9:6 and 6:6 complex experimental data with χ^2^ values of 2.22 and 24.5, respectively, whilst the SUN1-KASH5 6:6 crystal structure (PDB accession 6R2I; Gurusaran and Davies, 2021) fitted to the same 9:6 and 6:6 complex experimental data with χ^2^ values of 62.5 and 1.32, respectively.

### Protein sequence and structure analysis

Multiple sequence alignments were generated using *Jalview* (Waterhouse et al., 2009), and molecular structure images were generated using the *PyMOL* Molecular Graphics System, Version 2.0.4 Schrödinger, LLC.

### Statistics and Reproducibility

All biochemical and biophysical experiments were repeated at least three times with separately prepared recombinant protein material.

## Data availability

Crystallographic structure factors and atomic co-ordinates have been deposited in the Protein Data Bank (PDB) under accession numbers 7Z8Y, 8B46 and 8B5X, and corresponding raw diffraction images have been deposited at https://proteindiffraction.org/.

## Acknowledgements

We thank Diamond Light Source and the staff of beamlines I24 and B21 (proposal MX25233). This work was supported by a Wellcome Senior Research Fellowship to O.R.D. (Grant Number 219413/Z/19/Z), and a core grant to the Wellcome Centre for Cell Biology (203149).

## Author contributions

M.G. and B.S.E. crystallised SUN1-KASH6 and performed biophysical experiments. O.R.D. solved the SUN1-KASH6 crystal structures, analysed data, designed experiments and wrote the manuscript.

---

**Supplementary Figure 1:**
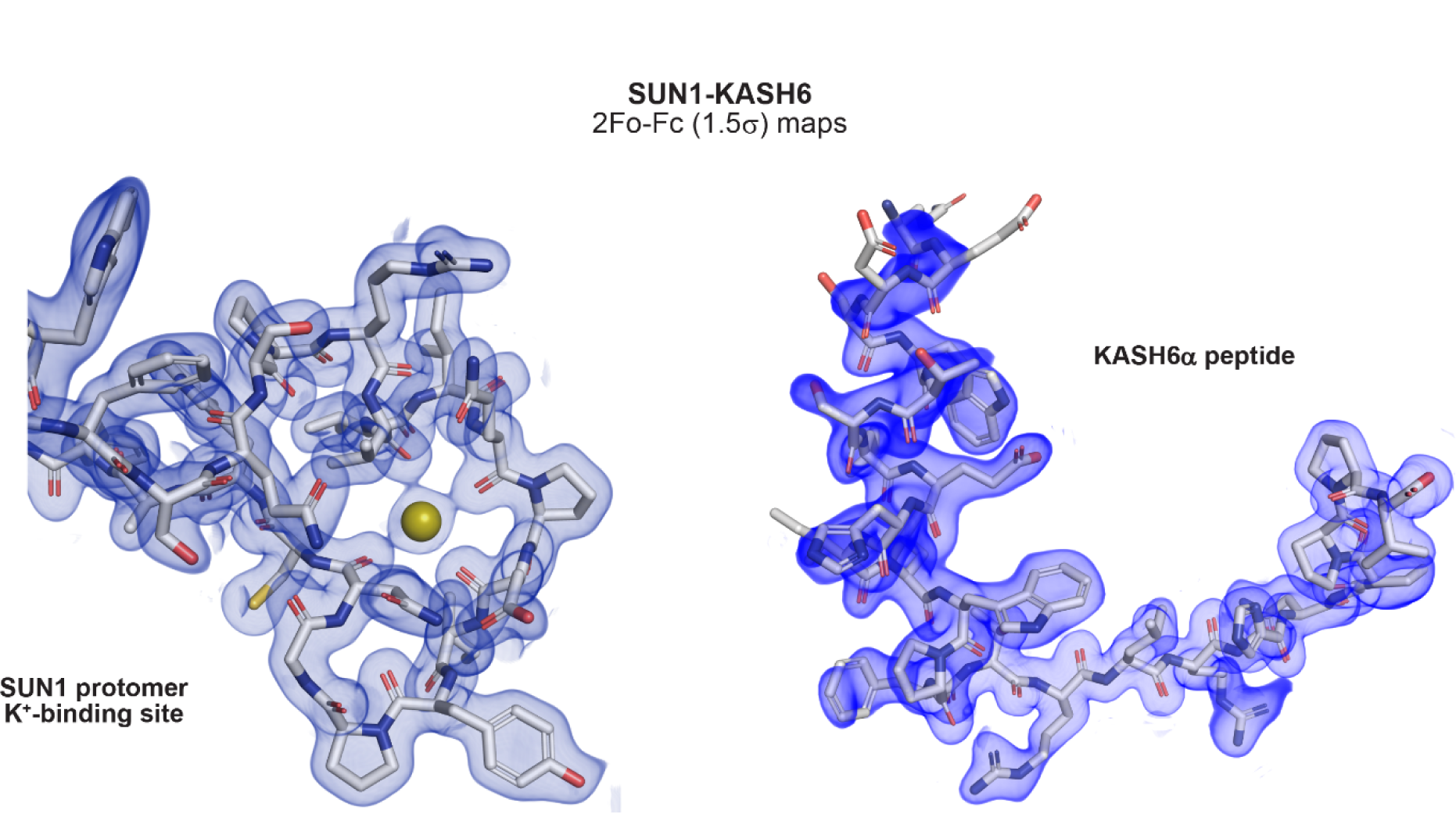
Crystal structure of SUN1-KASH6 (1.7 Å) in a 9:9 stoichiometry. 2Fo-Fc electron density maps (1.5σ) of SUN1-KASH6 (1.7 Å) for the potassium-binding site of the SUN1 protomer (left) and the KASH6α peptide (right).

**Supplementary Figure 2:**
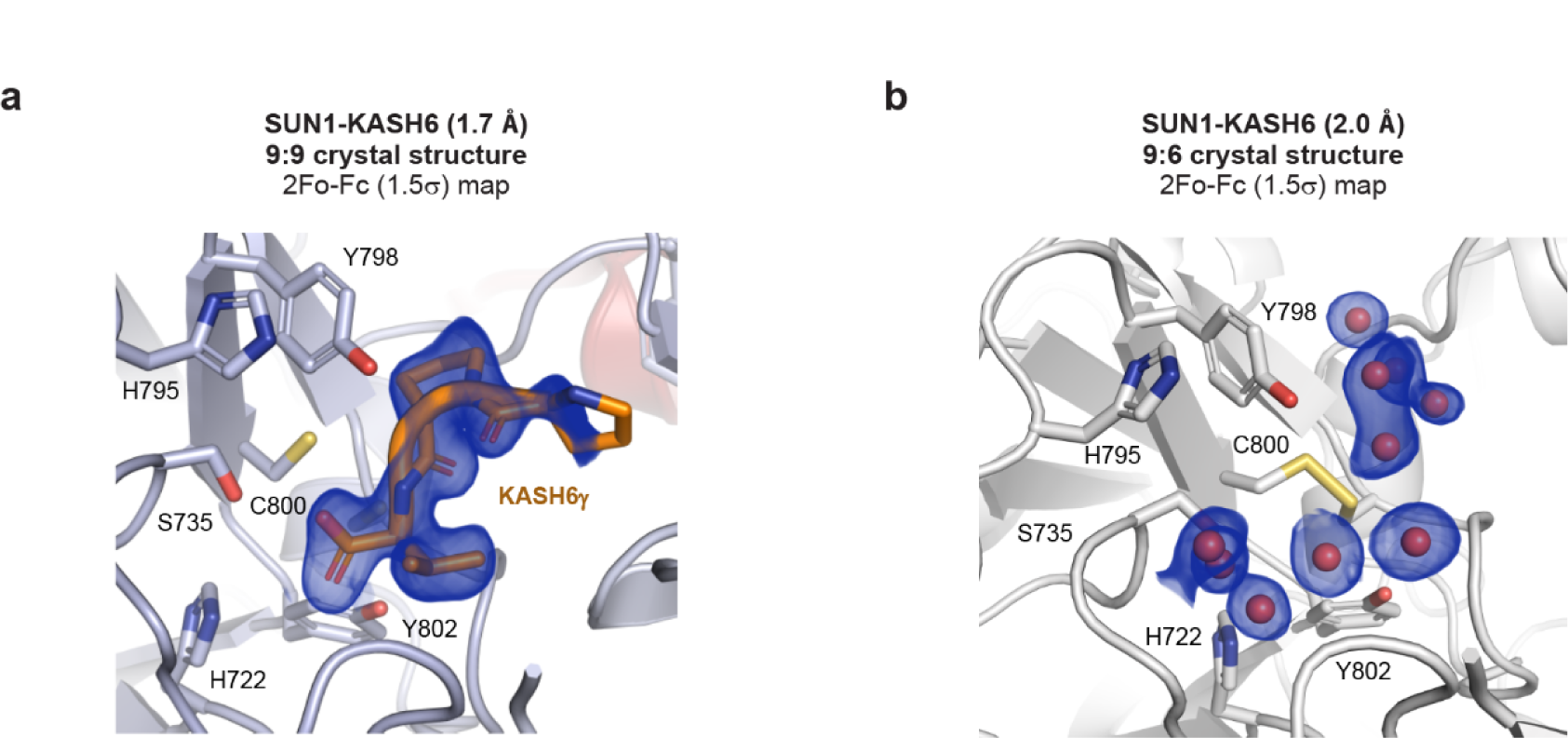
Crystal structures of SUN1-KASH6 at the KASH6γ-binding pocket. (**a,b**) 2Fo-Fc electron density maps (1.5σ) for the KASH6γ peptide and crystallographic water molecules at the KASH6γ-binding pockets of the (**a**) SUN1-KASH6 (1.7 Å) structure at 9:9 stoichiometry, and the (**b**) SUN1-KASH6 (2.0 Å) structure at 9:6 stoichiometry.

**Supplementary Figure 3:**
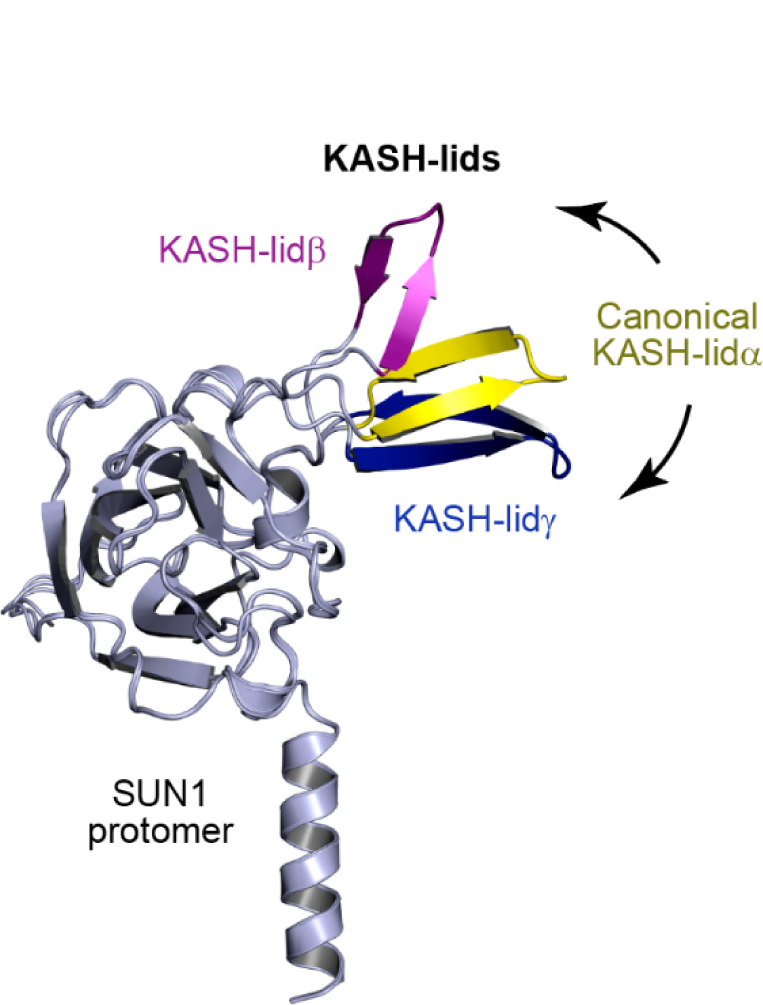
Alternative conformations of the three KASH-lids of the SUN1-KASH6 structure. Superposition of the three SUN1 protomers from one trimer of the structure, showing the canonical orientation of KASH-lidα (yellow), high angulation of KASH-lidβ (purple) and low angulation of KASH-lidγ (blue).

**Supplementary Figure 4:**
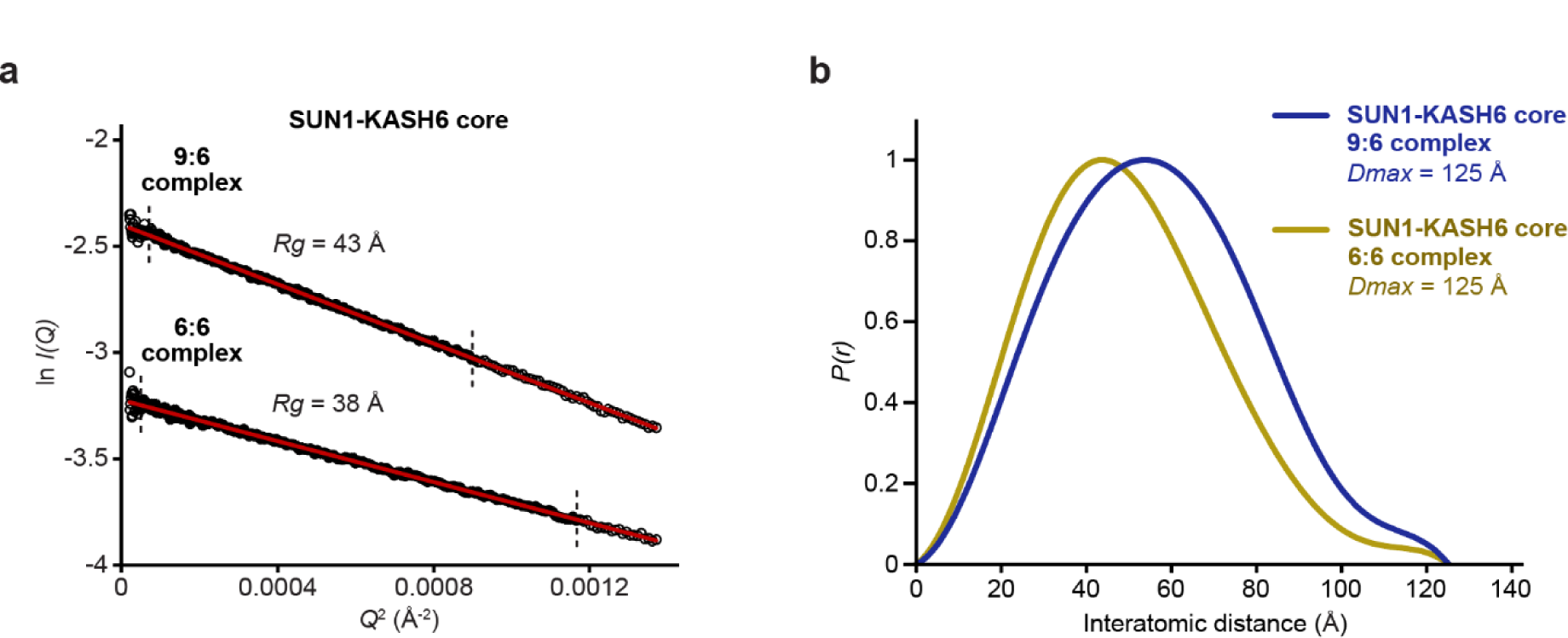
Small-angle X-ray scattering (SAXS) of SUN1-KASH6 core 9:6 and 6:6 species. (**a**) SAXS Guinier analysis to determine the radius of gyration (*Rg*); linear fits are shown in red, with the fitted data range highlighted in black and demarcated by dashed lines. The *Q*.*Rg* values were < 1.3 and *Rg* was calculated as 43 Å and 38 Å for 9:6 and 6:6 species, respectively. (**b**) SAXS *P(r)* interatomic distance distributions in which maximum dimensions (*Dmax*) were determined as 125 Å for both 9:6 and 6:6 species.

